# Induced illusory body ownership in Borderline Personality Disorder

**DOI:** 10.1101/628131

**Authors:** Eli S. Neustadter, Sarah K. Fineberg, Jacob Leavitt, Meagan M. Carr, Philip R. Corlett

**Author notes:** These authors contributed equally to this work (Eli S. Neustadter, Sarah K. Fineberg).

## Abstract

**Background:** One aspect of selfhood that may have relevance for Borderline Personality Disorder (BPD) is variation in sense of body ownership. We employed the rubber hand illusion (RHI) to manipulate sense of body ownership in BPD. We extended previous research on illusory body ownership in BPD by testing: 1) two illusion conditions: asynchronous & synchronous stimulation, 2) relationship between Illusion experience and BPD symptoms, and 3) relationship between illusion experience and maladaptive personality traits.

**Methods:** We measured illusion strength (questionnaire responses), proprioceptive drift (perceived shift in physical hand position), BPD symptoms (DIB-R score), and maladaptive personality traits (PID-5) in 24 BPD and 21 control participants.

**Results:** For subjective illusion strength, we found a main effect of group (BPD > HC, F = 11.94 *p* = 0.001), and condition (synchronous > asynchronous, F(1,43) = 22.80, *p* < 0.001). There was a group x condition interaction for proprioceptive drift (F(1,43) = 6.48, *p* = 0.015) such that people with BPD maintained illusion susceptibility in the asynchronous condition. Borderline symptom severity correlated with illusion strength within the BPD group, and this effect was specific to affective symptoms (r = 0.48, *p* < 0.01). Across all participants, trait psychoticism correlated with illusion strength (r = 0.44, *p* < 0.01).

**Conclusion:** People with BPD are more susceptible to illusory body ownership than controls. This is consistent with the clinical literature describing aberrant physical and emotional experience of self in BPD. A predictive-coding framework holds promise to develop testable mechanistic hypotheses for disrupted bodily self in BPD.

**Highlights:**

- The rubber hand illusion (RHI) allows measurement of self-disturbance.
- People with BPD had greater illusion susceptibility and this correlated with affective symptoms.
- Interoception stabilizes representations of body ownership, and is impaired in BPD.
- Illusion strength correlates with psychotic traits across levels of psychopathology.
- Predictive coding frameworks can probe mechanisms of impaired body ownership in psychopathology.

## 1. Introduction

### 1.1. The embodied self in Borderline Personality Disorder

#### 1.1.1. Self-disturbance is a core feature of BPD

Aberrations of self-experience and identity are considered a core symptoms of Borderline Personality Disorder (BPD) [1]. Self-disturbance is characterized by a markedly persistent unstable sense of self that can be realized by dramatic shifts in self-image, shifting goals and values, and feelings of emptiness, dissociation, and non-existence [2, 3]. These experiences are distressing and dangerous; in a qualitative study, Brown et al. [4] found that more than 50% of women with BPD and history of self-harm endorsed disturbances in self-experience, such as emptiness, numbness, or feeling dead, as reasons for non-suicidal self-injury.

#### 1.1.2. Bodily experience is disrupted in BPD

One aspect of selfhood that may have relevance for pathologies of self in BPD is the experience of body ownership. Indeed, abnormal bodily experiences in BPD are common, including bodily dissociation [5], altered pain perception [6], and deficits in interoception (the awareness and processing of internal bodily signals) [7].

Mechanistically, sense of body ownership is constituted by integration of sensorimotor (external) and interoceptive (internal) signals [8]. Neural computations on these signals generate a probabilistic, and therefore malleable, model of self-representation [9]. For a healthy person, sense of body ownership is stable and taken for granted, while in certain mental disorders such as BPD, sense of body ownership may be more variable and plastic. Experimental paradigms that directly manipulate the experience of body ownership have the potential to elucidate aberrations in embodied self-experience in BPD.

### 1.2. Probing the embodied self: The Rubber Hand Illusion

Illusions can test the plasticity of body ownership by manipulating integration of self and non-self stimuli. One paradigmatic body illusion is the Rubber Hand Illusion (RHI) [10]. During the task, a participant’s hidden hand is stroked in synchrony with an appropriately positioned and visible rubber hand (**Figure 1**). The RHI can induce the feelings that the rubber hand belongs to the participant (subjective illusion) and that the participant’s own hand has moved toward the rubber hand (proprioceptive drift). Typically, the RHI is measured by a self-report questionnaire of illusory experience (adapted from [10]) and the spatial magnitude of proprioceptive drift [11].

**Figure 1:**
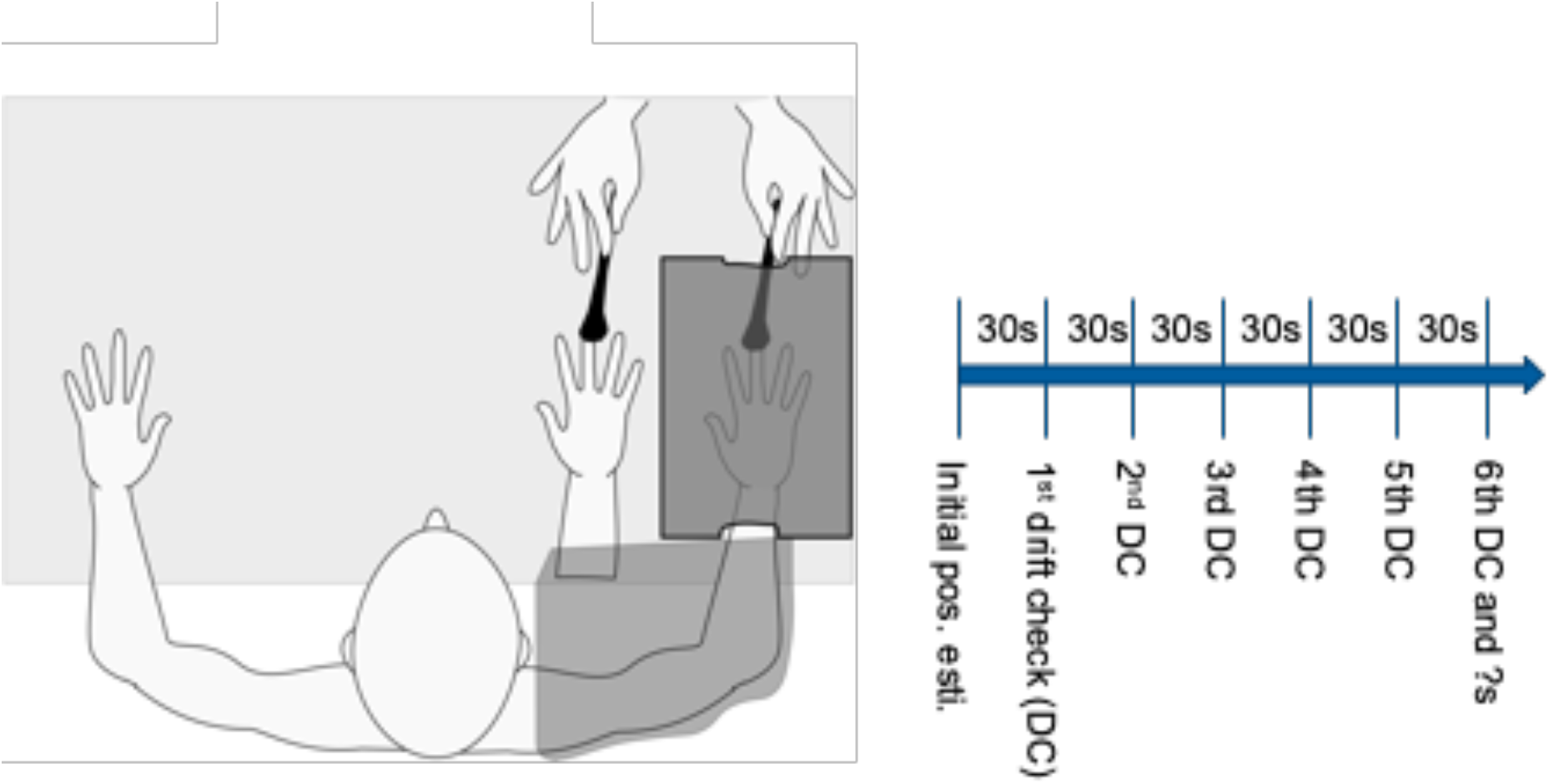
Rubber Hand Illusion setup. Participant’s gloved right hand is placed in cardboard box obstructing it from view. A life-like gloved rubber hand is placed medial to the box. A cloth is draped across right shoulder, covering the wrist of the rubber hand and the cardboard box proximally, where the participant’s real hand enters. During illusion induction, the participant is instructed to visually focus on the rubber hand while an experimenter provides brushstrokes to the middle phalanges of the real hand (through an opening in the cardboard box) and an equivalent location on the rubber hand for 3 minutes. Hand localization (“drift”) estimates are taken at 30 second intervals. The questionnaire is administered once after each stimulation condition (synchronous, asynchronous).

It is theorized that the RHI results from the multimodal (e.g. visuo-tactile) integration of sensory events in peri-personal space: an area including and immediately surrounding the body that is implicated in maintaining a dynamic cortical representation of the body [12]. RHI induction is sensitive to visuospatial plausibility and the timing of sensory stimulation, such that unrealistic placement of the rubber hand and temporally asynchronous stroking have been found to attenuate illusory body ownership in healthy participants [13, 14].

Eshkevari et al. [15] highlight two factors that promote induction of the rubber illusion. One factor, “visual capture,” occurs prior to visuo-tactile stimulation, whereby a sense of body ownership results from over-weighting of the visual stimulus of the rubber hand over proprioceptive information of the real hand. The other factor, which entails simultaneous seen and felt touch of the fake and real hand during simultaneous stroking, results in the illusion of rubber hand ownership via the multisensory integration of temporally co-occurring visual and tactile stimulation. Empirical data from healthy participants and computational modelling of rubber hand ownership demonstrate that the illusion can occur without tactile stimulation (first factor) and is enhanced by temporally synchronous (vs. asynchronous) stroking of fake and real hands [16]. Importantly, increased susceptibility to the first factor, which occurs in both synchronous and asynchronous conditions (as it occurs prior to tactile stimulation) may indicate imprecise bodily representations that result in the overweighting of exteroceptive information [16] [15].

### 1.3. Pathologies of illusory body ownership

The RHI has been conducted across a range of mental disorders in which anomalous self-experience has been implicated and which share clinical overlap with BPD, including schizophrenia [11], body dysmorphic disorder [17], and eating disorders [15, 18]. These conditions are associated with increased susceptibility to the RHI as measured by self-report questionnaire [18], proprioceptive drift [17] or both [11, 15]. Increases in subjective measures of the illusion and proprioceptive drift have also been demonstrated in pharmacological models of psychosis (i.e. ketamine) in healthy participants, implicating NMDA hypofunction and augmented neural oscillations in the gamma-range that promote cross-modal binding [13]. This interpretation was bolstered by the finding of maintained illusory experience in an asynchronous version of the RHI with pharmacologic challenge, highlighting the methodological importance of administering the task in both synchronous and asynchronous versions.

The vividness of the illusion has also been linked to schizotypy in healthy participants [19], suggesting that altered body ownership may be a marker of psychosis-proneness. However, the interpretation of these results is limited as task demand characteristics of the illusion questionnaire may not have been controlled for. In particular, the original Botvinick & Cohen [10] questionnaire was designed to include target and non-target items to control for suggestibility, but to our knowledge no clinical study has adequately assessed group differences in the relative endorsement of target and non-target items on the RHI questionnaire. Furthermore, target items (which probe the illusions of touch, causality, and ownership, respectively) sequentially probe qualitatively more encompassing aspects of the illusion. For example, people who minimally experience the illusion may only endorse the illusion of touch (feeling the touch on the location where the rubber hand is touched), while those who experience a stronger illusory experience may also endorse ownership of the rubber hand (that the rubber hand is <u>their</u> hand). However, differences in the relative endorsement of individual target items, both between and across clinical groups, has not been previously directly tested. Close examination of target item responses and attention to dimensional symptomology can reveal trait markers across diagnostic thresholds that may influence illusion susceptibility.

### 1.4. The current study: Probing body plasticity in BPD

To date, there has been one study examining illusory body ownership using the RHI in people with current and remitted BPD [20]. That study analyzed findings in a synchronous version of the task only, and found increased subjective experience of the illusion, but similar proprioceptive drift, in people with BPD compared to HCs. However, group differences in the relative endorsement of target and non-target items was not accounted for. Additionally, the authors found a small but significant correlation between illusory body ownership and state and trait dissociation after controlling for BPD symptom severity. We extend these findings by (1) testing group differences in the relative endorsement of target vs. non-target items, (2) probing responses to individual target items to explore more granular group differences in illusory experience, (3) directly testing hypotheses about asynchronous stimulation, and (4) exploring relationships between illusory body ownership and BPD symptom phenotypes and dimensional measures of maladaptive personality.

In the current study, we conducted the RHI task with people with BPD and healthy controls (HC) in temporally synchronous and asynchronous conditions. We made the following hypotheses:

**H1. Illusion strength would be greater in BPD vs HC groups in both synchronous and asynchronous conditions**

*We hypothesized that people with BPD would be more susceptible to the illusion as measured by both subjective questionnaire responses (H1.1) and proprioceptive drift (H1.2). This difference has been observed most strongly in synchronous condition in other settings; however, some increase in susceptibility in clinical groups has also been observed in the asynchronous condition. We are the first to report asynchronous condition results in BPD*.

**H2.1. Tactile illusion strength would be greater than ownership illusion strength.**

**H2.2. The illusion of ownership, but not the illusions of perception or causality, will be more strongly endorsed in BPD than in HC.**

*Some RHI studies descriptively report differential endorsement of target questions: Q1 (tactile illusion) versus Q2 (causality), Q3 (ownership), however these differences have not been directly tested. Given self-disturbance in BPD, we hypothesized that pair-wise comparisons of target-item endorsement would reveal specific increased endorsement of Q3 in BPD vs HC across conditions*.

**H3 (exploratory). Illusion strength would positively correlate to psychotic-like symptoms and traits.**

*In BPD, cognitive-perceptual disturbances are common, and psychotic-like traits are present in undiagnosed people in the general population. Given previous work linking RHI illusion strength to ketamine intoxication, psychosis, and schizotypy, we explored correlation of RHI illusion strength to psychotic-like experiences in BPD and HC, as well as BPD symptom clusters in BPD group*.

## 2. Methods

### 2.1. Subjects

This study was approved by the Yale Institutional Review Board. Results for this study were collected as part of a larger battery of experimental tasks. Results from those tasks as well as the recruitment strategy for these participants are described in detail elsewhere [21, 22]. Briefly, women aged 18-65 were recruited from the community. HCs had no current psychiatric conditions, and BPD participants had no current substance dependence and no primary psychotic disorder according to intake interview assessment (see **supplement 1**).

### 2.2. Symptom and self-report scales

HC and BPD participants completed a series of well validated self-report symptom scales and structured clinical interviews including: the Beck Anxiety Inventory (BAI) [23], Beck Depression Inventory (BDI-II) [24], The Personality Inventory for DSM-5 (PID-5) [25], The Structured Clinical Interview for DSM-IV Personality Questionnaire [26], and the Revised Diagnostic Interview for Borderlines (DIB-R) [27]). Please refer to **supplement 2** for information on scale validation and subscales.

### 2.3. Rubber hand illusion paradigm

Participants wore a non-latex glove on their right hand, sat in front of a table, and placed their right hand into an open cardboard box (**Figure 1**). All participants underwent RHI procedures on their right hand only as it was previously demonstrated that laterality and handedness had no effect on the subjective experience of the illusion or magnitude of proprioceptive drift, the two main outcomes measures for the task in the current study [28].

In the box, the participant’s right hand was occluded from their view, but not from the view of the experimenter who sat across the table facing the participant. A gloved life-sized rubber hand was positioned so that the hand was visible to the participant on the medial end of the box. A cloth was then draped over the participant’s shoulder covering both the real right arm and the arm of the rubber hand.

Before induction of the illusion, participants made an initial estimate of the spatial location on their occluded right hand via a numbered ruler that was placed on top of the box. Each participant then underwent synchronous and asynchronous versions of the task, each lasting 3 minutes. In the synchronous condition, an experimenter used the brush of a paintbrush to provide soft simultaneous touch at 1 Hz frequency in the proximal to distal direction along the middle phalanges of the real index finger and an equivalent location on the rubber hand. Procedures for the asynchronous condition were identical except that brush strokes were offset in time by 0.5 seconds (resulting in alternating touch on the real and rubber hands).

#### 2.3.1. Measure of subjective experience of the illusion

After synchronous and asynchronous conditions, participants completed a questionnaire adapted from Botvinick & Cohen [10] to assess their subjective experience of the illusion (**Table 1**). Variations of this questionnaire have been used widely in RHI research [17]. Similar to previous studies, the first three items (“target”) were used to create an index score as they are more strongly and consistently endorsed than the other items, and they reflect expected illusory experience [17]. The remaining items (“non-target”) have historically been included to control for suggestibility and task demand characteristics as they are endorsed only minimally by healthy samples [17]. However, they are often endorsed in clinical psychiatric populations and during pharmacologic challenge (e.g. ketamine) [13]. For each condition, a cumulative “target item” score was created as the average rating across items 1-3 and a cumulative “non-target item” score was created as the average rating across items 4-9. Significantly higher target scores compared to non-target scores was used as an indicator of successful induction of the illusion.

**Table 1:**
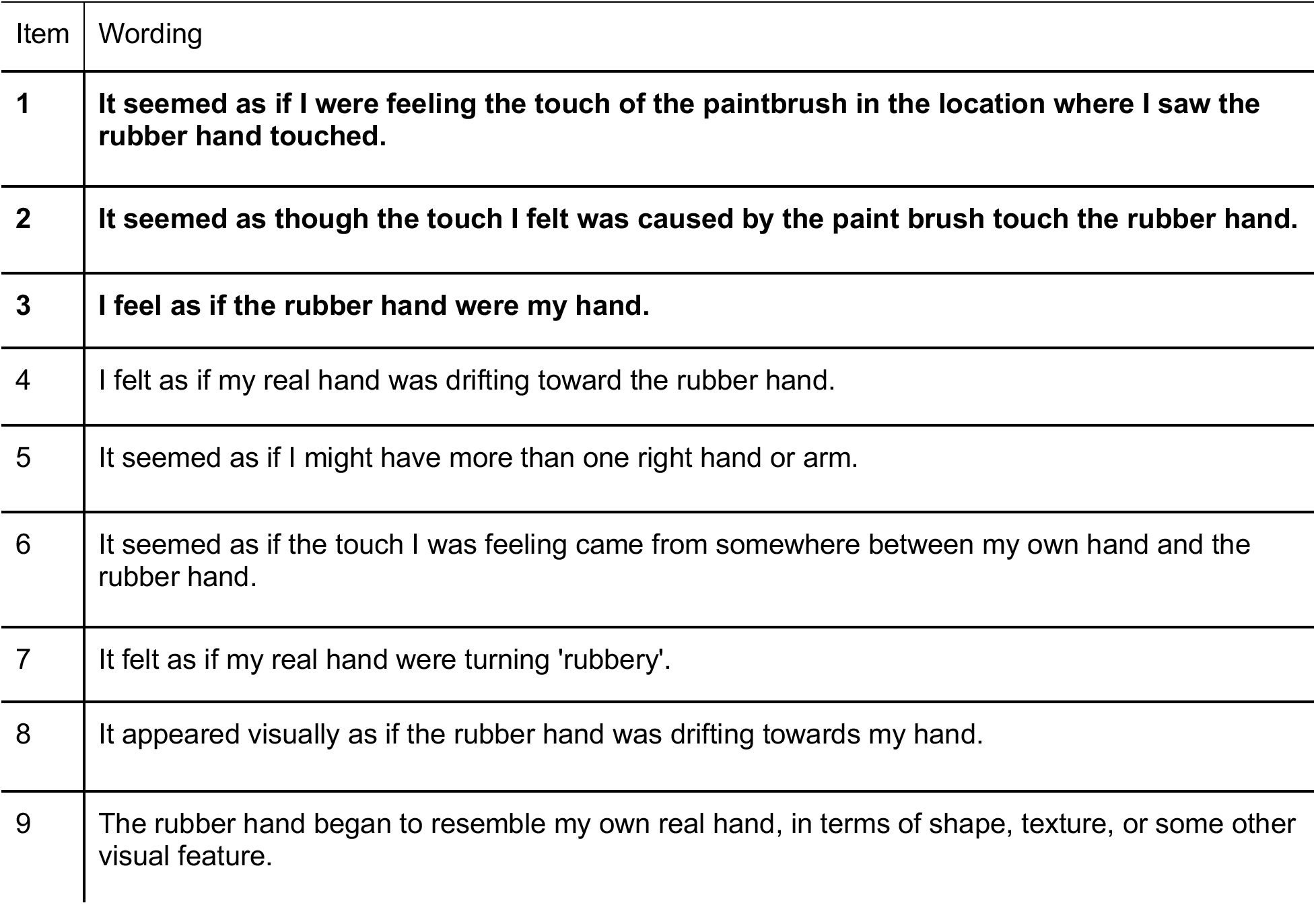
RHI Questionnaire Items (target items in bold)

#### 2.3.2. Measure of proprioceptive drift

Proprioceptive drift refers to the extent to which participants estimated their hand as being closer to the rubber hand after induction of the illusion. Participants estimated the position of their hidden right index finger before stimulation, and then at 30 second intervals during stimulations (6 times over 3 minutes of stimulation) by referring to a numbered ruler placed on top of the box. At each interval, participants were reminded not to move their hand. At each interval, the position of the ruler was jittered to prevent participants from anchoring on previous estimates [29]. Proprioceptive drift was calculated as the difference in estimated hand location between the pre-trial estimate and average of the six post-trial estimates. Positive values, then, refer to post-trial estimates that are closer to rubber hand than initial estimates. Positive drift values are consistent with successful induction of the illusion.

### 2.4. Planned statistical analyses

Parametric tests were conducted for analyses on main outcome variables (subjective experience questionnaire and proprioceptive drift) as values for skewness and kurtosis were all within −2 to 2, indicating normal univariate distribution [30].

To test for successful induction of the illusion in each group, we conducted a 2 x 2 analysis of variance ANOVA to compare the effects of condition (synchronous vs. asynchronous) and item-type (target vs. non-target) on subjective endorsement of the illusion.

Separate 2 x 2 ANOVAs were used to assess impact of group (HC vs. BPD) and condition (synchronous vs. asynchronous) on target item endorsement (hypothesis H1.1) and proprioceptive drift (hypothesis H1.2). ANOVAs were further explored with post-hoc t-tests. To assess for specificity of target versus non-target item endorsement, we performed a one-way ANCOVA to assess for group differences (HC vs. BPD) in target item endorsement using nontarget item endorsement as a covariate.

Repeated measures ANOVA test (2 group x 2 condition x 3 target items) was employed to test for differential endorsement of individual target items (hypothesis H2.1 and H2.2).

Lastly, we performed Pearson correlations to explore the relationships between RHI measures and symptom scales. Correlations were one-tailed unless stated otherwise to test for positive correlations (hypothesis H3).

Alpha values were set to 0.05 for primary analyses and, more conservatively, to 0.01 for post-hoc analyses and correlations. We report effect sizes using Cohen’s D for t-tests, and partial eta squared for ANOVAs.

## 3. Results

### 3.1. Participant characteristics

Twenty-four women were enrolled in the BPD group and 21 women were enrolled in the HC group. HC and BPD groups were matched on age, years of education, and race (**Table 2**). The BPD group was significantly more symptomatic on measures of BPD symptom severity (SCID-II, BSL, DIB-R), depression (BDI), and anxiety (BAI) (**Table 2**).

**Table 2:** Participant Characteristics. Mean results are reported followed by standard deviations in parentheses. Groups were matched on age, education, and race. All participants were female. BPD group participants reported significantly more anxiety, depression, and BPD symptoms than did HC participants.

|  | **BPD** | **HC** | **Statistics** |
| --- | --- | --- | --- |
| **n** | 24 | 21 |  |
| **Age (yrs)** | 33.17 (12.47) | 31.10 (13.13) | t = 0.54; p = 0.59 |
| **Education (yrs)** | 13.96 (2.46) | 15.02 (2.45) | t = -1.45; p = 0.15 |
| **<u>Race</u>** |  |  | Chi square = 3.19; df = 2; p = 0.74 |
| **Asian** | 8.30% | 9.50% |  |
| **Black** | 16.70% | 28.60% |  |
| **Hispanic** | 4.20% | 19% |  |
| **White** | 66.70% | 42.90% |  |
| **Not Reported** | 4.20% | 0% |  |
| **BAI** | 24.83 (12.80) | 7.35 (10.05) | t = 4.92; p < 0.001 |
| **BDI** | 23.04 (12.71) | 2.52 (4.52) | t = 7.39; p < 0.001 |
| **DIB-R (unscaled)** | 28.00 (6.55) | 3.19 (4.17) | t = 14.90; p < 0.001 |
| **SCID-II self report** | 9.75 (3.51) | 0.95 (1.40) | t = 11.31; p < 0.001 |
| **BSL-23** | 36.25 (21.42) | 3.95 (4.32) | t = 7.219; p < 0.001 |

### 3.2. Induction of illusory limb ownership

In both BPD and HC groups, target items were endorsed more strongly than non-target items in the synchronous condition. Furthermore, target items were more strongly endorsed in the synchronous condition compared to asynchronous condition (see **supplement 3** for statistics) Taken together, these results suggest that we were able to successfully induce the RHI in BPD and HC groups.

### 3.3. Self-Report RHI Questionnaire

*H1.1. Subjective illusion strength would be greater in BPD vs HC groups in both synchronous and asynchronous conditions*.

To test hypothesis H1.1, we tested for group differences in mean target item endorsement using a 2 group x 2 condition repeated measures ANOVA (**Figure 2A, 2B**). We found a significant main effect of group (BPD > HC, F(1,43) = 11.94, *p* = 0.001, η^2^ = 0.22) and of task condition (synchronous > asynchronous, F(1,43) = 22.80, *p* < 0.001, η^2^ = 0.35). No significant interaction was found (F(1,43) = 1.72, *p* = 0.681, η^2^ < 0.01).

**Figure 2:**
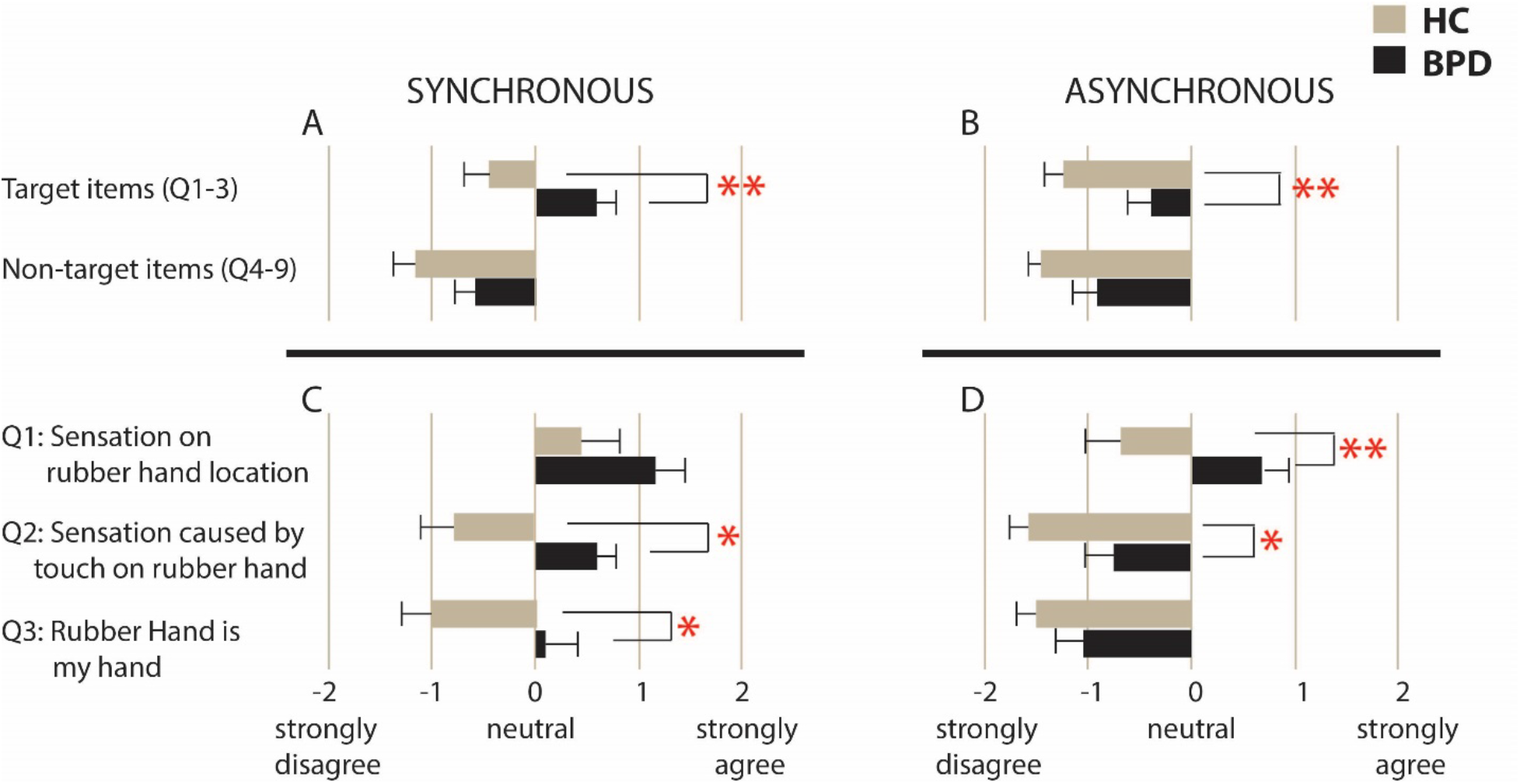
RHI Questionnaire Reponses. Averaged mean scores for target and nontarget items in synchronous (2A) and asynchronous conditions (2B). Error bars denote standard error of the mean. Means for individual target items are displayed for both synchronous (2C) and asynchronous (2D) conditions. * p < 0.05; ** p < 0.01

To determine whether group differences in target item endorsement in the illusion inducing condition (synchronous) could be accounted for by task demand characteristics or suggestibility, we conducted a one-way ANCOVA to test for a difference between BPD and HC groups in target item endorsement while controlling for responses to non-target items. We found that the effect of group remained significant when we controlled for the non-target items (F(1,42) = 4.40, p = 0.042, η^2^ < 0.1) suggesting that group differences in target item responses do reflect differences in the magnitude of illusory experience.

*H2.1. Tactile illusion strength would be greater than ownership illusion strength*.

*H2.2. Greater subjective illusion in BPD would be accounted for by ownership illusion*.

To test hypotheses H2.1 and H2.2, we conducted a repeated-measures ANOVA (2 groups x 2 conditions x 3 target items) (**Figure 2C, 2D**). We found a main effect of group (F(1,43) = 11.94, *p* = 0.001, η^2^ = 0.22), condition (F(1,43) = 22.80, *p* < 0.001, η^2^ = 0.35), and target item (F(1,43) = 26.16, *p* < 0.001, η^2^ = 0.38. While condition x group, item x group, and condition x item interactions were non-significant, we found a significant group x item x condition interaction (F(1,43) = 4.89, *p* = 0.032, η^2^ = 0.10). Post-hoc tests to unpack this 3-way interaction revealed that pair-wise comparison between Q1 and Q2 in the synchronous condition is significant in the control but not BPD group (control: t(20) = 3.12, *p* = 0.005, d = 0.68; BPD: t(23) = 1.75, *p* = 0.094, d =0.36). Other pairwise comparisons did not differ significantly by group.

To examine hypothesis H2.1, we conducted post-hoc t-tests comparing Q1 (tactile illusion) and Q3 (ownership illusion) in each condition. In all participants taken together, Q1 was more strongly endorsed than Q3 in both conditions (synchronous: t(44) = 4.17, *p* < 0.001, d = 0.63; asynchronous: t(44) = 4.79, *p* < 0.001, d = 0.72). Of note, Q2 and Q3 were endorsed comparably across conditions (Synchronous: t(44) = 1.61, *p* = 0.114, d = 0.24; Asynchronous: t(44) = 0.87, *p* = 0.39, d = 0.13). This suggests that the illusion of touch (i.e. feeling the touch on the rubber hand) was more easily induced by the RHI, while illusions of causality (i.e. felt touch was caused by touch on rubber hand) and ownership (“I felt as if the rubber hand were mine”) indicate more severe body illusion experiences that are more difficult to induce.

To examine hypothesis 2.2, we conducted post-hoc t-tests comparing group differences in individual target item endorsement. In the synchronous condition, BPD and HC groups comparably endorsed Q1 (t(43) = 1.59, *p* = 0.120, d = 0.48). Compared to HC, BPD endorsed Q2 (t(43) = 2.58, *p* = 0.013, d = 0.77) and Q3 more strongly (t(43) = 2.48, *p* = 0.017, d = 0.74); however, these differences did not achieve statistical significance at α = 0.01. In the asynchronous condition, BPD endorsed Q1 more strongly (t(43) = 2.77, *p* = 0.009, d = 0.83) compared to HC. Additionally, BPD endorsed Q2 more strongly, (t(43) = 2.34, *p* = 0.025, d = 0.70), but not at the statistical significance level of α = 0.01. Lastly, BPD and HC endorsed illusion of ownership at comparable levels t(43) = 1.21, *p* = 0.233, d = 0.36) in the asynchronous condition. In summary, contrary to our hypothesis, group differences in target item endorsement appear to be driven by different items in synchronous and asynchronous conditions. While in the synchronous condition group differences appear to be driven by differential endorsement of Q2 (illusion of causality) and Q3 (illusion of ownership), in the asynchronous condition, they are accounted for by differential endorsement of Q1 (tactile illusion) and, to a lesser extent, by Q2 (illusion of causality).

### 3.4. Proprioceptive Drift

*H1.2. Proprioceptive illusion strength would be greater in BPD vs HC groups in both synchronous and asynchronous conditions*.

To test hypothesis H1.2., we explored group differences in proprioceptive drift using 2 x 2 repeated measures ANOVA (group x condition) (**Figure 3**). Main effect of group (BPD vs HC, F(1,43) <0.001, *p* = 0.99, η^2^ < 0.01) and of task condition (synchronous vs asynchronous, F(1, 43) = 2.19 *p* = 0.15, η^2^ = 0.05) were not significant. However, a significant group x condition interaction was found (F(1,43) = 6.48, *p* = 0.015, η^2^ = 0.13). Post hoc paired sample T-tests demonstrated that, contrary to our hypothesis, while the HC group had significantly reduced proprioceptive drift in the asynchronous condition (t(20) = 2.90, *p* = 0.009, d = 0.63), the BPD group had no significant difference in drift across conditions (t(23) = 0.75, p = 0.462, d = 0.094) (**Figure 3**). We also found weak to moderate relationships between target item endorsement and proprioceptive drift that did not meet our significance cut-off of *p* < 0.01 (see **supplement 4**).

**Figure 3:**
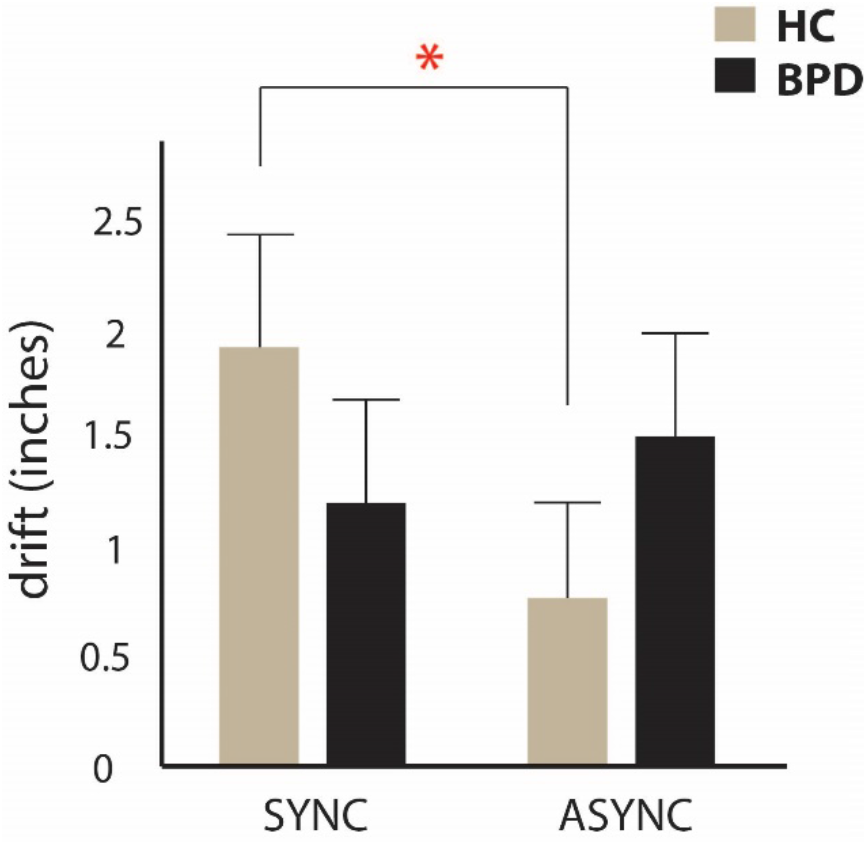
Proprioceptive drift. Mean drift toward rubber hand following six 30 second trials of synchronous (sync) or asynchronous (async) stimulation. Error bars represent standard error of the mean. * p < 0.05

### 3.5. Symptom/Trait Correlations

*H3. Exploratory: Illusion strength would positively correlate to psychotic-like symptoms and traits*.

#### 3.5.1. BPD symptoms

We investigated whether BPD symptom severity and BPD symptom clusters relate to illusion strength in the clinical group. To do so, we conducted one-tailed Pearson correlations between DIB total (unscaled score) and subscale (affect, cognition, impulsivity, and interpersonal relationship sections) scores (unscaled) and the following RHI measures: target-item score, item-3 (“I felt as if the rubber hand was my hand”) and proprioceptive drift in the synchronous condition (statistics in **Table 3**). We limited correlations to the synchronous condition to limit multiple comparisons and to focus on the more illusion-inducing condition. At the α = 0.01 level, we found that target-item index score and item-3 endorsement were related to the affect subscales with correlations in the large effect range. Proprioceptive drift was not related to BPD symptom severity or symptom clusters within the clinical group.

**Table 3:** Relationship between illusion susceptibility, BPD symptom clusters and maladaptive traits. The table displays Pearson correlations coefficients between proprioceptive drift, target-item endorsement, item three endorsement in the synchronous condition, and clinical/personality variables. On the right side are the scatterplots for the relationship between average target item endorsement in the synchronous condition and DIB affect in the BPD group (upper panel) and trait psychoticism as measured by PID-5 in the whole sample (lower panel). * p < 0.05, one tailed; ** p < 0.01, one tailed Note: DIB-R = Diagnostic Interview for Borderlines-revised. DIB-R includes affect, cognition, impulsivity, and interpersonal sub-scales. PID-5 = Personality Inventory for DSM-5 which has scales for the following maladaptive traits: negative affect, detachment, antagonism, disinhibition, and psychoticism. (s) Drift = proprioceptive drift in synchronous condition. (s) targ = average target-item response in synchronous condition. (s) Q3 = response to item 3 on RHI questionnaire in the synchronous condition: ‘’I feel as if the rubber hand were my own.”

| BPD<br>only | Scale | (s) Drift | (s) target | (s) Q3 |
| --- | --- | --- | --- | --- |
|  | DIB: Total score | 0.019 | .384* | .400* |
|  | DIB: Affect | 0.092 | .481** | .680** |
|  | DIB: Cognition | -0.289 | .411* | .358* |
|  | DIB: Impulsivity | 0.283 | 0.256 | 0.075 |
|  | DIB: Interpersonal | 0.095 | -0.041 | 0.107 |
| Total<br>sample | Scale | (s) Drift | (s) target | (s) Q3 |
|  | NegativeAffect | -0.172 | .309* | 0.183 |
|  | Detachment | 0.026 | .368* | .325* |
|  | Antagonism | .335* | 0.256 | 0.121 |
|  | Disinhibition | -0.067 | 0.228 | 0.173 |
|  | Psychoticism | 0.101 | .440** | .393** |

#### 3.5.2. Dimensional personality assessment

Next, we examined the relationship between RHI and dimensional measures of maladaptive personality traits across all participants. To do so, we conducted one-tailed Pearson correlations between PID-5 personality trait domains (negative affect, detachment, antagonism, disinhibition, and psychoticism) and the following RHI measures: target-item score, item-3 (“I felt as if the rubber hand was my hand”) and proprioceptive drift in the synchronous condition (**Table 3**). At the α = 0.01 level, only trait-psychoticism was significantly related to the target-item index score and item-3 endorsement, with correlations observed in the medium-effect range. Proprioceptive drift was not significantly related to clinical traits at the p < 0.01 level.

Of note, six participants (four BPD and two HC) did not complete the PID-5. The two HC participants were comparable to other HCs in age, education, BDI, BAI and RHI outcomes. The four BPD participants were both highly symptomatic and appeared to have higher target item endorsement in the synchronous condition. Thus, these results likely underestimate the correlation between maladaptive traits and subjective response to the illusion. Note that the very small sample size was prohibitive of inferential statistics.

## 4. Discussion

In this study, we have extended the previous investigation of illusory body ownership in BPD by directly assessing findings in the asynchronous condition, analyzing differential endorsement of self-report items, and identifying further associations with clinical and personality trait variables. In the paragraphs to follow, we will interpret RHI behavior in BPD within a predictive coding account of bodily self [31, 32], which posits that representations self are probabilistically generated through integration of top-down predictions about the body and bottom-up “prediction errors” of sensory inputs across interoceptive and exteroceptive domains.

We hypothesized that compared to HC, people with BPD would have greater target item endorsement (H1.1) and larger proprioceptive drift (H1.2) in both synchronous and asynchronous conditions. H1.1 was supported: BPD had greater target item endorsement in both conditions. Contrary to H1.2, we found a significant group x condition interaction on drift measurements: BPD and HC had comparable drift during synchronous stimulation. However, during asynchronous stimulation, BPD had maintained drift while HC had significantly reduced drift.

As hypothesized, we found increased body plasticity in BPD as measured by subjective endorsement of illusory experience. Bekrater-Bodmann et al. [20] reported increased subjective experience of the illusion; we clarified this finding by demonstrating that this group difference remained significant after controlling for the endorsement of non-target items, suggesting that increased target item response reflects alterations in the magnitude of illusory experience. We also extend their findings by demonstrating increased susceptibility in both synchronous *and asynchronous* conditions, indicating that illusion susceptibility occurs generally, rather than specifically during synchronous stimulation. While Bekrater-Bodmann et al. [20] employed asynchronous stimulation merely as a manipulation check, others have compared RHI results across conditions (e.g. [13, 15, 17]) to elucidate possible mechanisms underlying abnormalities in illusory body ownership. For example, Morgan et al. [13] found maintained illusory experience from synchronous to asynchronous stimulation during ketamine (an NMDA antagonist) challenge in healthy participants. NMDA antagonism (a model for early psychotic illness) is thought to weaken top-down signaling, leading to over-weighting of bottom-up input, even when the bottom-up signals are inconsistent. In the asynchronous RHI condition (a state of inconsistent bottom-up signals), the weaker top-down signaling produces a large prediction error regarding self-attribution, putatively resulting in illusion experience. In BPD, illusion susceptibility across synchronous and asynchronous conditions may similarly indicate weak top-down signaling regarding body-ownership.

RHI induction is hypothesized to arise from two processes [15]: 1) visual capture, which occurs prior to tactile stimulation, whereby rubber hand ownership is experienced via integration of visual and proprioceptive inputs of the fake and real hands, respectively; and 2), temporal integration of visual and tactile input during synchronous stroking. Studying RHI in eating disorders, Eshkevari et al. [15] interpreted maintained illusion susceptibility in asynchronous conditions as a heightened sensitivity to visual capture over distorted bodily signals. This interpretation was bolstered by the finding that interoceptive deficits were a significant predictors of illusory body ownership in ED. Importantly, interoception (i.e. the processing and awareness of internal bodily signals) is theorized as a central modality in stabilizing mental representations of bodily-self in predictive coding frameworks (e.g. [8, 32, 33]). Accordingly, the precision associated with prediction error of sensory input—the confidence or uncertainty ascribed to it— modulates the integration of bottom-up and top-down information flow, such that low precision-weighted prediction errors are less likely to update (top-down) prior beliefs. In the RHI, the stability of body ownership is maintained by the relative precision of interoceptive vs exteroceptive input. Reduced certainty, or “trustworthiness,” ascribed to interoceptive signals leads to the overweighting of exteroceptive input (e.g. seeing the rubber hand) during the task, resulting in increased susceptibility to the illusion (see [34] for empirical support). BPD is associated with deficits in interoceptive processing [35]. However, the relationship between interoception and body plasticity was not directly assessed in this study. Future research can assess the extent to which interoceptive processing, e.g. as measured by heart beat evoked potentials [35, 36], or heart beat detection [37], though see [38] for methodological limitations), mediates illusory body ownership in BPD and serves as a common mechanism of illusion susceptibility across personality, eating, and body-image disorders, for which there are symptomatic and clinical overlap [39, 40].

Contrary to H1.2, we found that BPD had comparable drift in both task conditions, while HC had significantly reduced drift in the asynchronous condition. While in previous studies, drift has been used as a “behavioral proxy” of rubber hand ownership (e.g. [41, 42]), Rohde, Di Luca and Ernst [43] found subjective endorsement of the illusion and drift to be separate and dissociable phenomena. In our sample, drift magnitude did not correlate with endorsement of RHI questionnaire items. Interestingly, Kaplan et al. [17] found that individuals with body dysmorphic disorder (BDD) demonstrate similar findings to our BPD sample such that they evidenced comparable drift in both conditions. They interpret this result in light of findings in healthy participants [43], that proprioceptive drift occurs to an equal extent during synchronous stroking and in a “just vision” condition (wherein participants estimate hand location after looking at rubber hand without tactile stimulation), while asynchronous stroking reduces drift by disrupting visio-proprioceptive integration. Kaplan et al. [17] posit that with regards to bodily awareness, people with BDD are less susceptible to the illusion-extinguishing effects of the asynchronous condition. If BPD shares a similar mechanism underlying maintained drift across conditions with BDD, this would be consistent with our proposed BPD self-model that is biased towards incorporating (even inconsistent) exteroceptive information in the setting of interoceptive deficits.

To our knowledge, differential endorsement of target-items has never been directly studied. Taking a closer look at responses to the RHI questionnaire, we hypothesized that the illusion of perception (Q1) would be more strongly endorsed than the illusion of ownership (Q3) (H2.1), and that greater subjective illusion strength in BPD would be accounted for by the illusion of ownership (H2.2). H2.1 was supported: across groups and conditions, Q1 (illusion of perception) was more strongly endorsed than Q3 (illusion of ownership). Contrary to H2.2., we found that group differences in target-item endorsement were driven by different questions in both conditions. H2.1 confirms our common-sense assumption that a tactile illusion is more easily inducible than the illusion of rubber hand ownership. Considering target item endorsement in the synchronous condition, we found comparable endorsement of Q1, but differential endorsement of Q2 and Q3 across groups, suggesting that similar perceptual experiences led to differential endorsement of statements regarding the relationship between real and rubber hands (i.e., that they are causally linked, or that the rubber hand is experienced as one’s own). Taken together, these findings are consistent with a predictive coding account of self-recognition [31], wherein more abstract multimodal self-representations are encoded at higher levels within a hierarchical model of self-processing. Intermediate-level beliefs are constrained by top-down expectations as well as sensory bottom-up information lower in the hierarchy. Thus, during the synchronous stroking, we hypothesize that the prediction error caused by RHI procedures could be accounted for at the level of a perceptual experience for healthy participants; whereas in BPD, RHI procedures lead to updating of more abstract self-representations, and therefore endorsement of causation and ownership illusions (Q2 and Q3, respectively). Similarly, while asynchronous stroking was sufficient to eradicate the illusion in HCs, the BPD group maintained an attenuated experience of the illusion (Q1 endorsement) related to perceptual experience.

Lastly, we performed exploratory correlations to assess the relationship between clinical traits and illusory experience. We hypothesized that psychotic-like experiences would be uniquely related to RHI illusion strength (H.3). In addition to linking illusion strength with psychotic-like experiences, we also found strong associations with affective symptoms in both the BPD group (as measured by the DIB affect subscale) and across the whole sample (as measured by PID-5 trait negative affect). While the link between psychotic-spectrum experience and RHI has been demonstrated in other settings (e.g. [19] [13] [13]), we are the first to demonstrate this association within BPD, providing further evidence that body plasticity may track psychosis-proneness trans-diagnostically. Finding a link between dissociation and RHI susceptibility in BPD, Bekrater-Bodman et al. [20] posit that altered NMDA neurotransmission may underlie altered body plasticity in the condition. This suggestion is bolstered by neurochemical evidence [44] implicating glutamatergic signaling in the anterior cingulate cortex (ACC) in BPD. Importantly, the ACC and the insular cortex have been identified as central structures for interoception [35]. Computational perspectives on mood and emotion suggest that emotional states reflect the certainty (or precision) regarding the interoceptive consequences of action, such that negative emotion “contextualize events that induce expectations of unpredictability” ([45], p. 2278). Thus, state negative affect may contribute to overweighting of exteroceptive input during RHI procedures in the setting of low precision-weighted interoceptive input. Clarifying the contribution of state, e.g. affect, and trait, e.g. emotion regulation [35], characteristics to the plasticity of body ownership also may elucidate the relationship between emotion arousal and clinical states such dissociation and depersonalization associated with BPD [5]. Alterations in body plasticity may also inform our understanding of interpersonal difficulties in the condition. BPD is associated with a two-fold increase in preferred interpersonal distance in live dyadic contexts compared to healthy controls, suggesting alteration in embodied peri-personal space [21]. Given the theoretical links between interoception, emotion, and theory of mind [46], targeting body awareness (e.g. through mindfulness practice [47, 48]) may be an important focus, especially for people with BPD whose symptom profiles are high in self-disturbance and psychoticism.

The correlation we observe between illusion susceptibility and psychotic-like traits here points at another mechanistic path through predictive coding to the observed results. We have written elsewhere about the critical role of priors in explanation-making in the face of a chaotic environment. For example, people with early psychosis suffer a barrage of difficult to explain perceptual experiences, likely related to aberrant signaling of prediction errors. We and others have demonstrated that top-down suppression of aberrant prediction error is a mechanism of odd belief formation, especially delusions in psychosis and psychotic-like states [49–52]. Strong priors for predictable causality may, by this logic, drive acceptance of the rubber hand as one’s own hand (as a way of explaining the conflicting visual and tactile cues). This may serve as a common mechanism of illusory body ownership across a wide variety of people with psychotic-like traits.

The findings from this work are best understood within the context of several limitations. Sample size was small, and subjects were all women. The small sample size prohibits examination of the impacts of potentially interesting demographics (race, sexual orientation, age) and co-morbidities on outcome. Inclusion of only biologically female, female-identified subjects for the study was important to decrease potential sources of variability in results given the small sample, but does limit generalizability of results. From a task set-up perspective, we did not include a baseline acclimation period to test for illusion induction from visual stimulus alone prior to tactile stimulation. This has been done in a non-clinical sample [16] and would enable the direct assessment of the relative contribution of visual capture vs. integration of visuotactile stimulation in producing enhanced illusory experience in clinical population. Furthermore, in a computational model, Majed, Chung, & Shams [16], demonstrated that the perception of body ownership as measured by the RHI can be described as a Bayesian causal inference. Future research applying this modeling techniques to clinical data can further probe to what extent increased body plasticity in BPD is driven by weakening of top-down representations of body-schemas, strong priors for rubber hand ownership, and/or bottom-up integration of interoceptive and exteroceptive input.

## Acknowledgements

We would like to thank Ada Umuego for her help creating figures. We also thank our research participants.

## Data availability

Data will be made readily available upon request to the corresponding author.

