## Supplemental Material for "Induced illusory body ownership in Borderline Personality Disorder"

**S1: Methods- Recruitment and enrollment**

Women aged 18-65 were recruited from the community and local clinics through flyers and online advertisements. Initial screening was performed over telephone using the BPD section of the Diagnostic Interview for Personality Disorders [1] and questions to assess for exclusion criteria. Based on results of the screen, participants were invited to the laboratory and informed consent was obtained. Participants were then screened in person using the Structured Interview for DSM-IV [2], and the Revised Diagnostic Interview for Borderlines (DIB-R) [3]. To be enrolled in the study, HCs had no current psychiatric conditions, and BPD participants had no current substance dependence and no primary psychotic disorder. On DIB-R, enrolled HCs scored ≤ 4 (scaled total), and enrolled BPD participants scored ≥ 8 (scaled total). Enrolled participants had no history of neurologic injury or illness. We also collected information on participant self-reported race and education level.

**S2: Symptom and self-report scale**

The Beck Anxiety Inventory (BAI) is a 21-item self report scale that has established reliability (Cronbach's alpha 0.94) and validity (r = 0.54 versus diary reports of anxiety) in an initial psychometric validation study [4, 5]. In our sample, 20 HC subjects and 23 BPD subjects completed the scale.

The Beck Depression Inventory (BDI-II) is a 21-item scale with established reliability (Cronbach's alpha 0.9) and validity (r = 0.71 – 0.86 versus a range of commonly used depression scales) in large meta-analysis [6]. In our sample, 21 HC subjects and 23 BPD subjects completed the scale.

The Personality Inventory for DSM-5 (PID-5) is a 220 item self-rated personality trait scale that assesses personality trait facets. The facets can be grouped into 5 broader trait domains: Negative Affect, Detachment, Antagonism, Disinhibition, and Psychoticism. Domain scales have established reliability (Cronbach's alpha ranging from 0.84-0.96 for trait domains) and validity (r = 0.44-0.67; median r = 0.53 for each train domain compared to conceptual counterpart of the Personality Psychopathology Five [7, 8]. In our sample, 18 HC subjects and 20 BPD subjects completed all trait domain scales. 1 HC subject completed subscales for antagonism, disinhibition, and psychoticism only.

The Structured Clinical Interview for DSM-IV Personality Questionnaire was not initially designed as a stand-alone instrument, but it has been found to have few false-negatives, and to be reliable compared to the clinician-administered version [9] [10]. In our sample, 21 HC subjects and 24 BPD subjects completed this questionnaire.

The Revised Diagnostic Interview for Borderlines (DIB-R) [3], is a diagnostic interview used to diagnose BPD with established internal reliability (e.g. Cronbach's alpha = 0.82 in an adolescent sample) [11], and validity (for cutoff of scaled score of 8, the DIB-R had a sensitivity of 0.82, a specificity of 0.80, compared to clinician-assessed DSM-III diagnostic criteria) [3]. The interview consists of 186 standard questions and 108 scores. Responses are totaled to score 22 descriptive statements reflecting features of BPD. Scores are then totaled for diagnostic determination and symptom burden assessment. Symptom burden is additionally characterized into 4 symptom-cluster subscales: affect, cognition, impulsivity, and interpersonal relationships. In our sample, 21 HC subjects and 24 BPD subjects completed this questionnaire.

**S3: Results- Induction of illusory limb ownership**

For each group (HC and BPD), we conducted a 2 x 2 ANOVA test to compare the effects of condition (synchronous/asynchronous) and item-type (target/non-target) on subjective endorsement of the item. We found a main effect of condition in both groups, with greater endorsement after synchronous than asynchronous stimulation (**BPD:** $F(1,23) = 12.59$, $p < 0.005$, mean score sync = 2.98, SE 0.19, mean score async = 2.31, SE 0.20, **HC:** $F(1,20) = 11.87$, $p < 0.05$, mean score sync = 2.25, SE = 0.20, mean score async = 1.67, SE 0.17), .  We also found a main effect of question type with greater endorsement of target versus non-target items (**BPD**: $F(1,23) = 17.60$, $p < 0.001$, mean score target = 3.02, SE = 0.18, mean score non-target = 2.27, SE 0.20, **HC:** $F(1,20) = 11.13$, $p < 0.005$), mean score target = 2.16, SE = 0.19, mean score non-target = 1.75, SE 0.16. Taken together, these results suggest that we were able to successfully induce subjective experience of the RHI in both BPD and HC groups. Also, the synchronous condition is more illusion-inducing than the asynchronous condition, and the target items are more strongly endorsed than the non-target items.

**S4: Results- Relationship between proprioceptive drift and RHI questionnaire**

Across groups, there were weak correlations that fell short of statistical significance between proprioceptive drift and target item endorsement in the synchronous ($r = 0.20$, $p = 0.099$, one-tailed) and asynchronous ($r = 0.22$, $p = 0.072$, one-tailed) conditions. While this relationship was not significant in the HC group, there was a moderate relationship with trend-level significance in the synchronous condition ($r = 0.32$, $p = 0.066$, one-tailed) and a moderate statistically significant relationship in the asynchronous condition ($r = 0.45$, $p = 0.013$, one-tailed) in the BPD group.
